# Compartmental structure in the secondary lymphoid tissue can slow down *in vivo* HIV-1 evolution in the presence of strong CTL responses

**DOI:** 10.1101/2024.06.24.600465

**Authors:** Wen-Jian Chung, Dominik Wodarz

## Abstract

Human immunodeficiency virus (HIV-1) replicates in the secondary lymphoid tissues, which are characterized by complex compartmental structures. While Cytotoxic T lymphocytes (CTL) readily access infected cells in the extrafollicular compartments, they do not home to follicular compartments, which thus represent an immune-privileged site. Using mathematical models, we have previously shown that this compartmental tissue structure can delay the emergence of CTL escape mutants. Here, we show computationally that the compartmental structure can have an impact on the evolution of advantageous mutants that are not related to CTL recognition: (i) Compartmental structure can influence the fixation probability of an advantageous mutant, with weakened selection occurring if CTL responses are of intermediate strength. (ii) Compartmental structure is predicted to reduce the rate mutant generation, which becomes more pronounced for stronger CTL responses. (iii) Compartmental structure is predicted to slow down the overall rate of mutant invasion, with the effect becoming more pronounced for stronger CTL responses. Altogether, this work shows that *in vivo* virus evolution proceeds slower in models with compartmental structure compared to models that assume equivalent virus load in the absence of compartmental structure, especially for strong CTL-mediated virus control. This has implications for understanding the rate of disease progression.

## Introduction

Human immunodeficiency virus infection is characterized by an initial acute phase, followed by a prolonged asymptomatic or chronic phase, and finally culminates in the development of AIDS. While the processes that drive the disease from the asymptomatic phase to AIDS are still incompletely understood, viral evolution is thought to be an important driver of disease progression [1, 2]. HIV mutates with a relatively fast rate [3, 4], resulting in beneficial mutations that allow the virus to replicate more efficiently, change aspects of cell tropism, and escape from immune responses [5, 6]. These viral evolutionary processes have been studied in detail, both in clinical and experimental settings [1, 7–9], and with the help of mathematical models [10–17]. Mathematical models are informed by experimental data, including data on virus load and estimates of viral parameters and immune cell kinetics. Most of these mathematical models consider the population of HIV-infected cells as a well-mixed system [18, 19]. At the same time, however, the bulk of virus replication occurs in the secondary lymphoid tissue, which consists of compartments with different characteristics. In particular, data suggest that it is important to distinguish between the extrafollicular (EF) and the follicular (F) compartments [20–31]. While virus replication occurs in both compartments, anti-viral cytotoxic T lymphocyte (CTL) responses can efficiently fight the infection in the extrafollicular, but not the follicular compartments. The follicles represent an immune-privileged site because of the limited ability of CTL to home to the follicles. Hence, in the presence of strong CTL responses, there is a marked discrepancy in the amount of virus replication occurring in the two compartments, with low virus load in the EF compartment and relatively high levels of viral replication occurring in follicular T helper cells. This has been shown in experiments with simian immunodeficiency virus (SIV)-infected macaques [22, 23, 32]. The discrepancy between virus load in the F and EF compartments was relatively large in animals characterized by more efficient overall virus control, while a more equal distribution of virus in the compartments was observed in animals with weaker CTL responses or during the acute phase of the infection before SIV-specific CTL responses have emerged. In SIV-infected macaques, the discrepancy in the amount of virus replication in the two compartments is especially prominent in elite controllers, where productive virus is largely cleared by CTL in the EF compartment, while replication continues to occur largely undisturbed by CTL in the follicles [22, 30]. The persistent productive virus replication in the follicles of elite controllers raises concerns about the ability of the virus to evolve, which can contribute to the loss of the elite controller status.

When these compartmental dynamics are taken into account in mathematical models, we have previously shown that the emergence of CTL escape mutants can be substantially delayed in a compartmental system, compared to a non-compartmental system with equivalent virus load [33]. This might contribute to explaining the observed unexpectedly long times until CTL escape mutant emergence in people living with HIV (PLWH) during chronic infection [34]. The reason is the lack of substantial selection for CTL escape mutants in the follicular compartments, where CTL are largely absent. Uncertainty, however, remains about how the compartmental structure of the secondary lymphoid tissues influences the evolutionary dynamics of other advantageous mutants that do not cause escape from CTL responses, such as mutants that replicate faster or infect cells more efficiently. Now selection forces are likely identical in both compartments. While a subdivision of a population into compartments has been shown to not affect the fixation probability of advantageous mutants in constant populations (depending on assumptions) [35–37], it has been shown to increase the time until mutant fixation [38, 39]. In addition to a population being subdivided into compartments, however, those compartments can have different population sizes, which might further influence evolutionary dynamics [40]. Moreover, these dynamics might be more complex if the variation in population size are driven by variation in the presence of immune responses (CTL). In the current study, we address this gap by investigating how the compartmental structure in the secondary lymphoid tissues, and the CTL-privileged nature of the follicles, influences the evolutionary dynamics of advantageous viral mutants that are unrelated to CTL escape. We consider several measures of virus mutant evolution, including the probability of mutant fixation, the rate of mutant generation, and the average time until mutant invasion in the presence of mutation processes.

### The mathematical model

We build on our previous mathematical models that described virus-CTL dynamics in the follicular and extrafollicular compartments of the lymphoid tissue [33, 41]. Hence, we take into account uninfected cells, cells infected by wild-type (WT) virus, cells infected by mutant virus, and CTL in the extrafollicular compartment (denoted by X_e_, Y_1e_, Y_2e_, and Z_e_, respectively), as well as equivalent populations in the follicular compartment (denoted by X_f_, Y_1f_, Y_2f_, and Z_f_, respectively). The dynamics can be described by the following set of ordinary differential equations, which track the development of these populations over time.

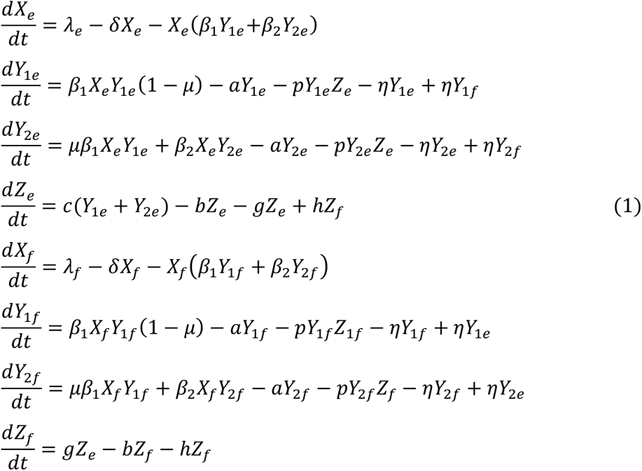

The virus dynamics are the same in both compartments. Susceptible target cells are produced with rates *λ_e_* and *<u>λ</u>_f_*, die with rates *δX_e_* and *δX_f_*, and become infected by WT and mutant viruses with a rate *X_e_(β_1_Y_1e_+β_2_Y_2e_)* and *X_f_(β_1_Y_1f_+β_2_Y_2f_)*. Upon infection of a susceptible cell with WT virus, the mutant is generated with a probability *μ*. Infected cells are assumed to die with rates *aY_ie_* and *aY_if_*, and are further killed by CTL with rates *pY_ie_Z_e_* and *pY_if_Z_f_*. Exchange of infected cells between the two compartments occurs with a rate *η*. CTL in the EF compartment expand in response to stimulation by both viral strains with a rate *c(Y_1e_+Y_2e_)*, die with a rate *bZ_e_*, and move to the follicular compartment with a rate *gZ_e_* (which is assumed to be low). In the follicular compartment, CTL do not expand. They may die with a rate *bZ_f_*, and home back to the EF compartment with a rate *hZ_f_* (assumed to be large). Note, that in our formulation the rate of CTL expansion is not proportional to the number of CTL, as in some previous studies [19], because this results in less oscillatory dynamics that are biologically more realistic. Free virus is not taken into account explicitly. Instead, the free virus population is assumed to be in quasi-steady state with the infected cell populations [19].

In the absence of evolution (*μ=0*), this type of model has been mathematically analyzed in our previous publication [33], and we refer the reader to this study for further details, which specifies basic properties such as conditions for the successful establishment of infection, and the dependence of virus load on CTL parameters. Here we assume that the virus establishes a persistent infection in both compartments, and that the amount of virus replication is sufficient to stimulate an ongoing CTL response. In the absence of mutations, the system then converges to a steady state in which the populations of uninfected cell, infected cells, and CTL persist in both the F and the EF compartments [33]. Relevant for our current work, the number of infected cells at equilibrium in the EF compartment is inversely proportional to the rate of CTL expansion (which we also refer to CTL strength, given by parameter *c*). Thus, the stronger the CTL response, the lower virus load in the EF compartment. Under the assumption of low infected cell migration rates and a low movement rate of CTL form the EF to the F compartment, the equilibrium number of infected cells in the follicular compartment is not substantially influenced by the strength of the CTL response. This recreates the experimental observation that the equilibrium number of infected cells in the two compartments is similar for weak CTL responses, but increasingly discrepant for stronger CTL responses (Figure S1) [22].

To determine the effect of compartmentalization on mutant evolution, we need to compare results from model (1) to a control model without compartmentalization but identical virus load. For a given parameter combination, we thus consider a one-compartment version of model (1) that is summarized in the Supplementary Materials section 2, but adjust the strength of the CTL response (parameter *c*) such that the equilibrium number of WT-infected cells in the one-compartment model is identical to the total equilibrium number of WT-infected cells in the two-compartment model (*Y_1e_+Y_1f_*), in the absence of mutants. Identical infected cell populations sizes are needed for comparison because differences in population size alone (rather than differences arising from compartmentalization per se) can affect the evolutionary dynamics of mutants.

Regarding parameters, we assume that the death rate of infected cells is given by *a=0.45*, consistent with experimental data [42], and other host/viral parameters are chosen arbitrarily such that the basic reproductive ratio of the virus is around the experimentally estimated average of R_0_≈8 [43]. Since clinical data display a spread of estimated R_0_ values, we will also repeat the analysis for a lower and higher value of R_0_. The mutation rate of HIV has been estimated to be on the order of 10^-5^ per base pair per generation [3], although more recent studies argue for a higher physiological rate [4]. The mutation rate, however, has to be considered in the context of the infected cell population sizes, and the total number of infected cells in the follicular and extrafollicular compartments are unknown. We therefore start with an arbitrary target cell population size such that the infection persists in the context of stochastic simulations. We then repeat the analysis by varying population sizes and mutation rates. There is uncertainty regarding some of the compartmental kinetics, such as the rate of virus movement from one compartment to the other (*η*), or the rate of CTL migration from the EF to the F compartment (*g*) and vice versa (*h*). From the experimental data summarized in the Introduction section, we know that the rate of virus exchange between compartments (*η*) must be relatively small, and that the rate of CTL migration from the EF to the F compartment (*g*) must also be small. If this was not the case, the behavior of the model would converge to that of a mixed system with similar virus population sizes in the two compartments even in the presence of a strong CTL response, which is inconsistent with experimental data [22]. However, a range of low *η* and *g* values are consistent with biologically realistic compartmental dynamics (see Figure S1). We will start with arbitrarily chosen low migration rates that qualitatively reproduce the kind of compartmentalization seen in experiments, and subsequently repeat simulations with different movement rates that still yield significant compartmentalization.

In our studies of virus mutant evolution, we performed stochastic simulations of ODE model (1) by using the Gillespie algorithm [44]. In these simulations, populations approach the equilibrium outcomes of the ODEs and stochastically fluctuate around these levels. In the absence of mutants, this is shown in Figure 1 for the two-compartment model and the one-compartment control model, for different CTL strengths (parameter c). For the two-compartment model, this shows an increased discrepancy in virus loads between the F and EF compartments for stronger CTL responses, as seen in experimental data [22]. For the weakest CTL response (lowest value of *c*), the infected cell population size in the two compartments is similar (Figure 1A, top panel). For stronger CTL responses, the number of infected cells in the EF compartment declines and shows stronger fluctuations around equilibrium, caused by CTL-mediated activity (Figure 1, top panel). Virus load in the F compartment and the degree of population fluctuations remain largely unaffected by changes in the parameter *c*, due to the very limited CTL activity in this location (Figure 1, top panel). For the one-compartment control model, virus load decreases, and population fluctuations increase for stronger CTL responses (Figure 1, bottom panels).

**Figure 1.**
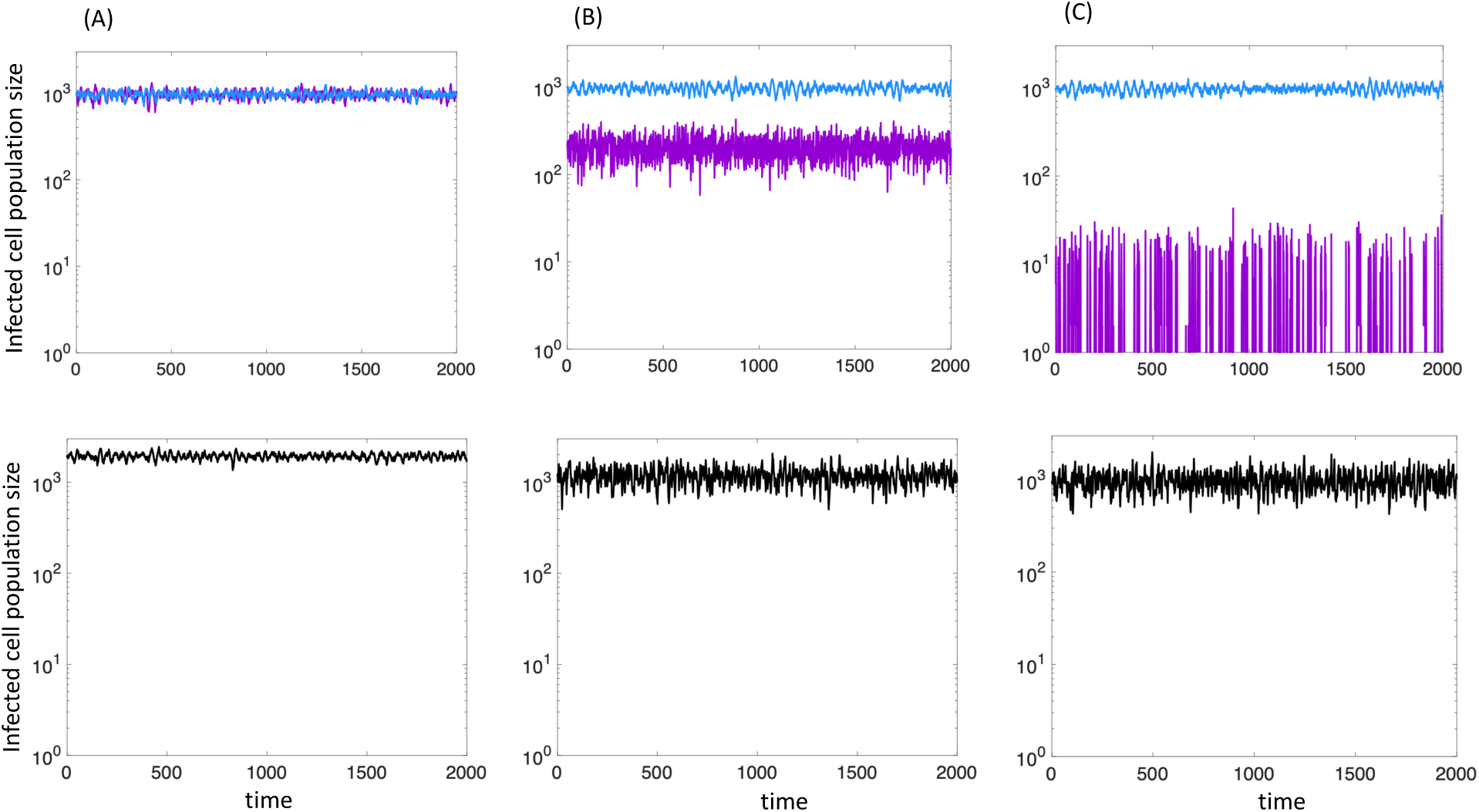
Gillespie simulations of model (1) with two compartments (top panels) and the one-compartment control model (bottom panels) for different CTL response strengths, expressed by variation in the parameter *c*. The infected cell population size in the F and EF compartments are shown in blue and purple colors, respectively. (A) Weak CTL response, *c=0.*0001 for the two-compartment model. (B) Intermediate CTL response, *c=0.1* for the two-compartment model. (C) Strong CTL response, *c=10* for the two-compartment model. In each case, the value of the parameter c in the one-compartment control model has been adjusted such that the equilibrium number of infected cells is identical to the total number of infected cells in the two-compartment model, as described in the text. Parameters were chosen as follows: λ_f_=500; λ_e_=500; β_1_=0.00007; δ=0.01; a=0.45; p=0.05; b=1; η=0.0001; g=0.0001; h=1.

We consider an advantageous mutant, expressed by a higher rate of infection, i.e. *β_2_>β_1_*. The advantage is identical in the two compartments. Therefore, the competition dynamics between the wild-type and the mutant virus strain are the same as in a single compartment model, which is well understood in the literature [19]. That is, the advantageous mutant grows and replaces the wild-type virus over time. The focus of our investigation here is to define how the compartmental structure influences the mutant invasion dynamics.

Different evolutionary measures will be analyzed here: (i) We start by investigating how compartmental structure influences the fixation probability of a mutant virus that is placed into a wild-type virus population at equilibrium, and how it affects the conditional time to fixation, i.e. the average fixation time in instances where the mutant does successfully invade. (ii) Next we examine how the compartmental structure affects the rate at which mutant viruses are generated. (iii) Finally, we put these processes together and determine the effect of compartmental structure on the average time to mutant fixation in a setting where the wild-type virus population replicates at equilibrium and mutates.

### Mutant fixation probabilities: compartmental structure can weaken selection

The fixation probability is determined by placing one mutant-infected cell into the wild-type-infected cell population at equilibrium, and numerically recording the fraction of simulation instances in which the mutant reaches fixation (rather than going extinct). Mutant fixation probabilities are well-understood in constant population models [45], such as the Moran or Fisher Wright processes, and a subdivision of the population into equally sized demes has not been found to alter mutant fixation probabilities under assumptions that are applicable here [35–37]. Subdivision of populations into demes of unequal sizes, however, has been found to weaken selection [45], and predator-prey type population fluctuations have been shown to also weaken selection [46]. Since both aspects apply to our current scenario, we investigated the fixation probability of an advantageous mutant in our model.

In our simulations, a single mutant-infected cell was introduced into either the F or the EF compartment with a probability given by the proportion of infected cells in each location. Figure 2A shows the relative mutant fixation probabilities for varying strengths of the CTL response (given by variation in the parameter *c*), assuming a 1% fitness advantage of the mutant. This is defined as the mutant fixation probability determined numerically in our computer simulations divided by the mutant fixation probability calculated from the constant population Moran process in an undivided population, given by 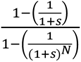, where *s* denotes the fitness advantage of the mutant and *N* the number of wild-type-infected cells at equilibrium. The mutant fixation probability displays a non-monotonic dependence on the CTL response strength for the 2-comparmtent model, with a minimum at an intermediate CTL responsiveness. For weak CTL responses (low *c*, when the infected cell population sizes in the EF and F compartments are similar) and for strong CTL responses (large c, when most infected cells are located in the F compartment), the observed mutant fixation probability is close to that predicted by the Moran process (indicated by a relative fixation probability close to 1). For intermediate CTL strength, the observed mutant fixation probability deviates most from the Moran process prediction, indicating weakened selection. In contrast, the mutant fixation probability declines monotonically for the 1-compartment control model, showing increasingly weakened selection for stronger CTL responses (Figure 2A).

**Figure 2.**
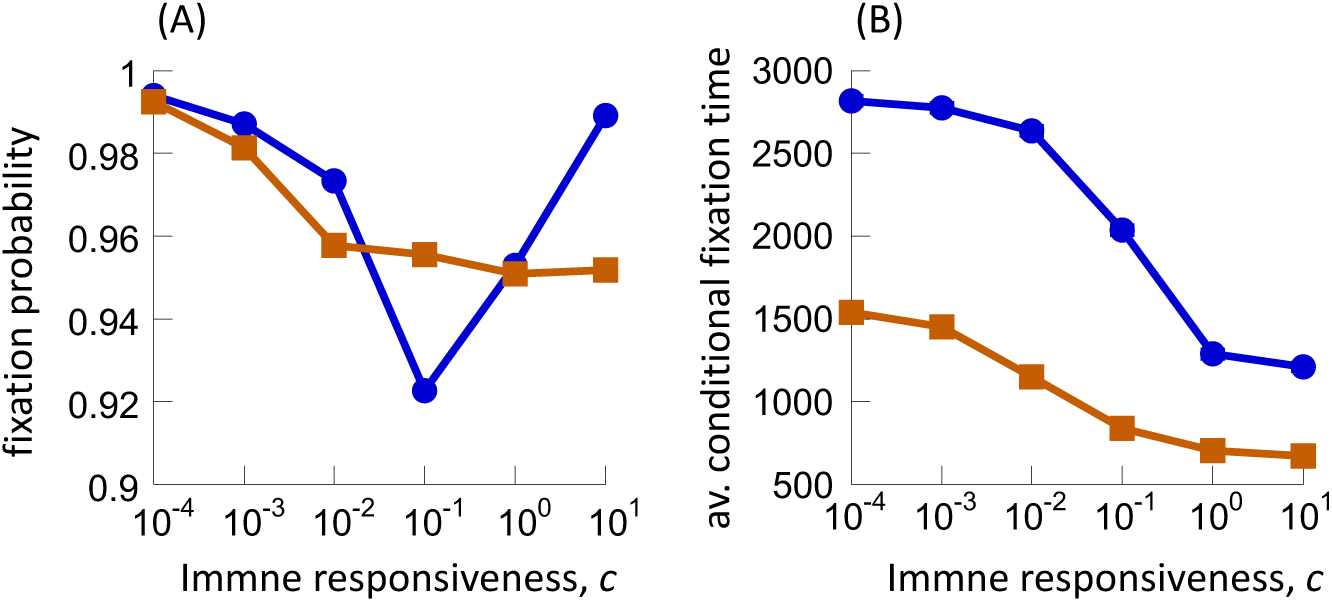
(A) Fixation probability and (B) average conditional fixation time of a mutant with a 1% advantage as a function of the immune responsiveness, c. Results for the two-compartment model are shown in blue, and those for the one-compartment control model in brown. Fixation probabilities were obtained computationally based on 10,120,000 simulation repeats. For the average conditional fixation times, standard errors are plotted but are too small to see. The average times are based on 115,000 simulation repeats. Parameters were chosen as follows: λ_f_=500; λ_e_=500; β_1_=0.00007; β_2_=1.01xβ_1_; δ=0.01; a=0.45; p=0.05; b=1; η=0.0001; g=0.0001; h=1.

Figure 2B plots the conditional fixation times of mutants, i.e. the average time it takes the mutant to reach fixation in those realizations that do result in mutant fixation. For both the two-compartment and the one-compartment control model, conditional fixation times decline with stronger CTL responses. Conditional fixation times are larger for the two-compartment model compared to the one-compartment control model (Figure 2B).

Two forces apply to our system that can in principle account for the weakened selection observed in our simulations (compared to standard Moran process predictions). These are population fluctuations brought about by the CTL response (predator-prey dynamics, [46]), and the population structure involving two demes with unequal population sizes [47]. In the one-compartment control model, CTL-induced fluctuations in the infected cell population appears to drive the reduced mutant fixation probabilities for stronger CTL responses (Figure 1). While CTL-induced fluctuations in the infected cell population also occur in the two-compartment model in the extra-follicular locations and can contribute to weakened selection, it appears that the discrepancy in infected cell population size between the F and EF compartments plays a major role in determining how the mutant fixation probability changes with the strength of the CTL response. To investigate this, we simulated a single-species two-patch constant population death-birth Moran process with migration between the patches, where the population sizes in the two patches corresponded to the number of infected cells at equilibrium in the EF and F compartment in model (1) for the various values of parameter *c*. (Supplementary Materials, section 3). This represents a much simpler scenario (single population, no infection dynamics), where the distribution of population sizes across the two patches is identical to those in our infection model, in the absence of any population fluctuations. As can be seen in Supplementary Figure S2A, the dependence of the relative fixation probability of an advantageous mutant on patch population sizes in the Moran model is qualitatively identical to the one observed in our two-compartment infection model, showing a minimum and weakened selection for population sizes that correspond to an intermediate CTL strength in model (1).

Therefore, this pattern is primarily a result of unequal population sizes among the two patches, similar to results reported by [47], and not due to CTL-induced population fluctuations. The weakened selection is entirely due to mutants that are placed into the smaller (corresponding to the EF) compartment, see Supplementary Figure S2B. The reason is that the influx of wild-type individuals from the larger to the smaller compartment reduces the advantage of the mutant in the smaller compartment. With this in mind, the trend for the two-compartment model in Figure 2 can be explained as follows: For weak CTL responses, the relative mutant fixation probability is high and similar to the Moran prediction because the infected cell population sizes in the two compartments is almost identical. Although for very strong CTL responses, the discrepancy in the infected cell population size between the compartments is highest, it is very unlikely that the mutant is placed in the EF compartment with the small population size. Hence, the relative fixation probability is again high and close to the Moran prediction. For an intermediate CTL strength, there is a sufficient difference in the infected cell population size between the two compartments, yet the population size in the EF compartment is still sufficient such that an initial mutant placement there is likely. This accounts for the reduction in the overall fixation probability of the mutant in the two-compartment model.

Section 4 in the Supplementary Information explores how these patterns can depend on the extent of the mutant fitness advantage and the infected cell population size. Selection is weakened to a lesser degree for larger mutant fitness advantages, and for larger overall infected cell population sizes.

### Compartmental structure reduces the rate of virus mutant generation

Now we introduce mutant generation by wild-type virus into the simulation We start with the wild-type virus-infected cell population at equilibrium and record the time at which the mutant is generated for the first time in the Gillespie simulations (Figure 3A). For more efficient CTL-mediated virus control (larger value of c), the time to first mutant generation increases, both in the 2-compartment model and in the 1-compartment control model. This increase, however, is substantially more pronounced in the 2-compartment model compared to the 1-compartment control.

**Figure 3.**
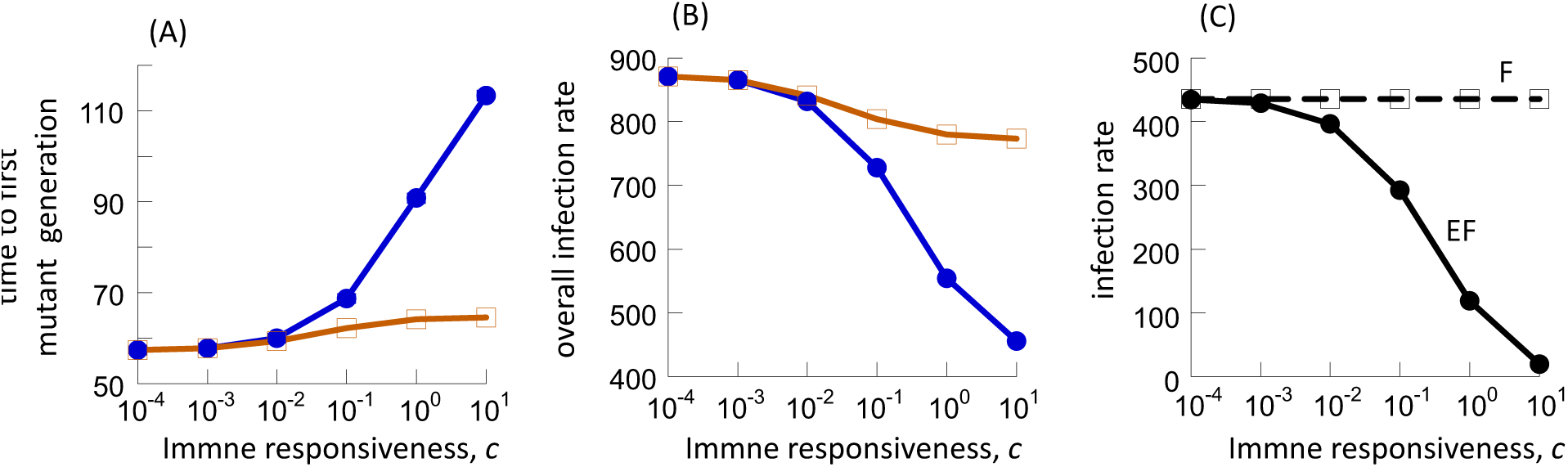
The rate of mutant generation. (A) Average time to first mutant generation determined by Gillespie simulations, depending on the CTL responsiveness, c. Standard errors are plotted but too small to see. Results are based on 115,000 simulation runs for each value of *c*. Blue and brown colors show results from the two-compartment and one-compartment control models, respectively. (B) The infection rate in the ODEs. (C) The infection rate for the two-compartment model in the ODEs, plotted separately for the F and EF compartments. Parameters are the same as in Fig 2. The mutation rate was *μ=2×10^-5^*.

This is also reflected in the rate of mutant generation in ODE system (1), which is proportional to the rate of infection, i.e. *β*_1_*X*_*e*_*Y*_1*e*_ and *β*_1_*X*_*f*_*Y*_1*f*_(Figure 3B). This measure declines for stronger CTL responses, and this decline more substantial in the two-compartment compared to the one-compartment control model, explaining the trend seen in panel (A). The reasons for these findings are as follows. In the follicular compartment, the population sizes of infected and uninfected cells are relatively stable across different CTL response strengths, *c*, because the CTL have limited access to the follicles. Hence the parameter *c* does not substantially influence the rate of mutant generation there (Figure 3C). In the extra-follicular compartment, however, an increase in the CTL responsiveness, *c*, leads to a reduction in the equilibrium number of infected cells, which is proportional to *1/c*. However, it only leads to an asymptotic rise in the availability of uninfected target cells (which approaches *λ/d*). Since the product of these two measures determines the infection rate (i.e. the rate of mutant generation), this approaches zero in the EF compartment for strong CTL responses (Figure 3C). In the 1-comparmtent control model, on the other hand, the equilibrium number of infected cells is assumed to be equal to the total number of infected cells across both compartments in the two-compartment model. Hence, there is no situation in which the number of infected cells approaches very low values (essentially zero), while the number of uninfected cells asymptotically approaches *λ/d*. Consequently, for increasing CTL strength, the total rate of infection at equilibrium (and hence the rate of mutant generation) is higher in the one-compartment control model compared to the two-compartment model.

### Compartmentalization delays the overall time to mutant fixation

So far, we considered the fixation probability and conditional fixation times of a mutant that is placed into a wild-type population at equilibrium (without the occurrence of continuous mutational processes). We also investigated the rate of mutant generation assuming that the wild-type virus population at equilibrium produces mutants during the process of infection. Here, we put both aspects together. We start with a wild-type-infected cell population at equilibrium and allow mutants to be generated with a rate *μ* upon infection. The simulation is run until the mutant replaces the wild-type population in both compartments. The average time to mutant fixation is determined based on repeated realizations of the computer simulation. We again consider a 1% advantage of the mutant, expressed by a higher rate of infection. Figure 4 shows the average mutant fixation time in the two-compartment model as a function of the CTL response strength, *c*, and this is again compared to the one-compartment control model with equal virus load. We start by considering a mutation rate that is low relative to the population size of infected cells (*μ=2×10^-5^).* For all values of *c*, we find that the average mutant fixation time is longer for the two-compartment model compared to the one-compartment control (Figure 4A). For weaker CTL responses, this difference is relatively small. The discrepancy grows for stronger CTL responses. The average mutant fixation time rises substantially for stronger CTL responses in the two-compartment model, while the increase is relatively small in the one-compartment control model (Figure 4A). Hence, in the presence of strong CTL responses, the compartmental structure of the secondary lymphoid tissue is predicted to substantially delay the time until advantageous mutants rise, compared to a scenario without compartmental structure but identical virus load.

**Figure 4.**
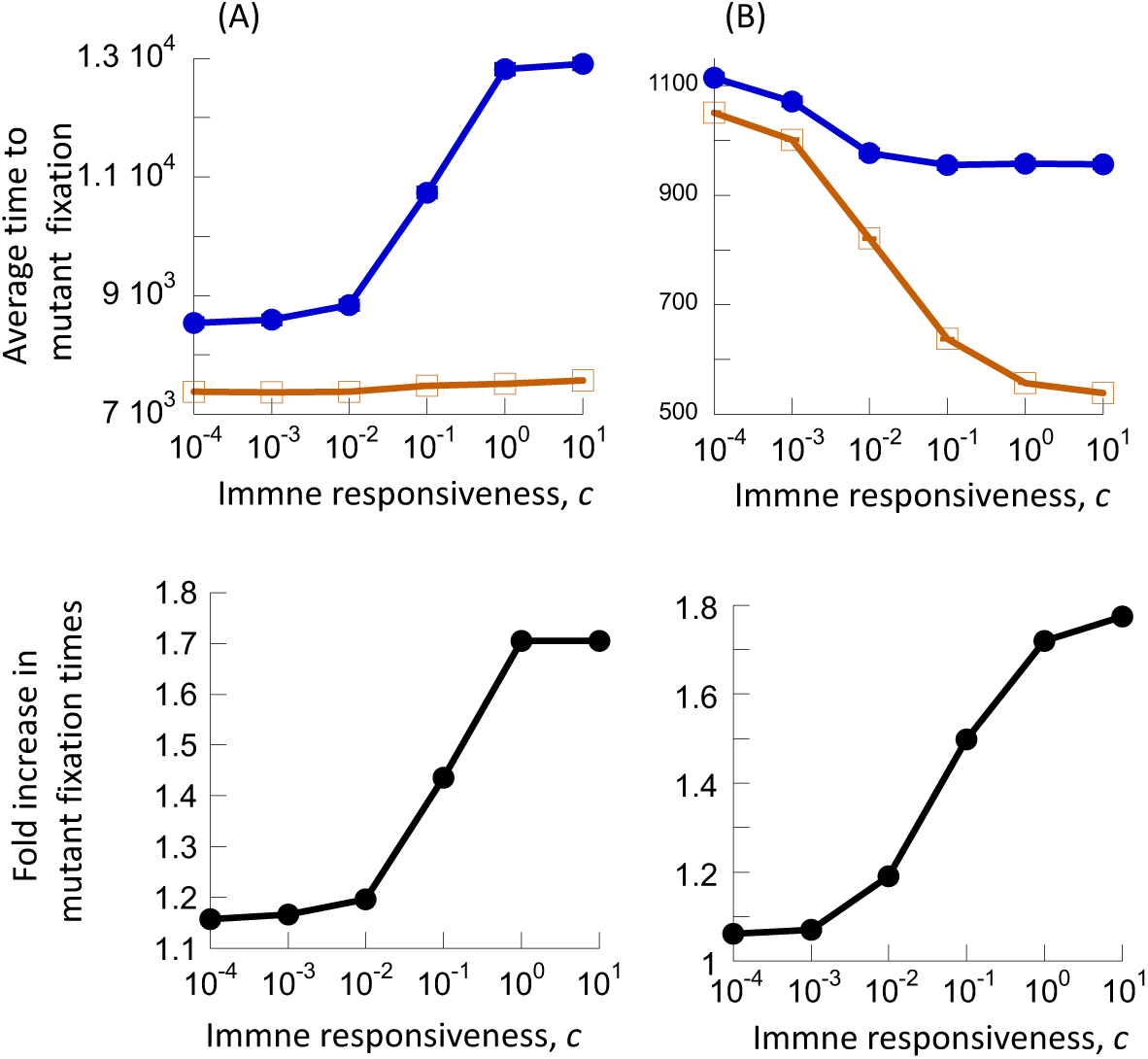
Average time to mutant fixation in the presence of mutation processes, based on Gillespie simulations of model (1). (A) Relatively low mutation rate, *μ=2×10^-5^*. (B) Higher mutation rate, *μ=2×10^-3^*. Top graphs show the average time to mutant fixation in the two-compartment model (blue) and the one-compartment control model (brown). Standard errors are plotted but too small to see. Bottom graphs show the fold increase of mutant fixation time in the two-compartment model compared to the one-compartment control model. Remaining parameter values are the same as in Fig 2. Results are based on 115,000 simulation runs for each value of *c*.

The substantial increase of the average mutant fixation time with stronger CTL responses observed in the two-compartment model can be counter-intuitive because stronger CTL responses lead to lower equilibrium numbers of infected cells, which should reduce the time to mutant fixation (Figure 2). Due to the low mutation rate relative to the infected cell population size, however, the generation of mutants is the rate-limiting step here, and this measure declines substantially with stronger CTL responses in the two-compartment model (Figure 3). This causes an increasing delay in the timing of mutant fixation for higher values of *c* in the two-compartment model.

If the mutation rate is high relative to the infected cell population size, the compartmental structure of the secondary lymphoid tissue is also predicted to delay the time for advantageous mutants to rise compared to the one-compartment control model with identical virus load. In Figure 4B, we see that that the average time to fixation is always longer in the two-compartment compared to the one-compartment control model, and this discrepancy again grows for stronger CTL responses. In contrast to the results observed for lower mutation rates, however, the average times to mutant fixation decline for stronger CTL responses, both for the two-compartment model and the one-compartment control model. The reason is that now mutant generation is not the rate-limiting step anymore (due to the faster mutation rate). Instead, the time of mutant invasion following mutant generation is the rate limiting step, and this becomes shorter for stronger CTL responses because of overall lower infected cell population sizes (Figure 2). The effect of compartmentalization, however, remains identical to the simulations with lower mutation rates: compartmentalization delays mutant invasion compared to the one-compartment control model with the same virus load, and this effect becomes more pronounced for stronger CTL responses (Figure 3B, bottom panel).

These dynamics are also shown with individual realizations of the simulated evolutionary dynamics (Figure 5). Panel A shows the dynamics for relatively low mutation rates, both for weak CTL-mediated control (Figure 5Ai) and for strong CTL-mediated control corresponding to elite control (Figure 5Aii). This shows that mutant production is a limiting step, and that this takes longer for stronger CTL-mediated virus control. Figure 5B repeats these simulations for relatively large mutation rates, again showing weak (Figure 5Bi) and strong (Figure 5Bii) CTL activity. This demonstrates that mutants are produced immediately and the rate of mutant growth is the main determinant of mutant invasion.

**Figure 5.**
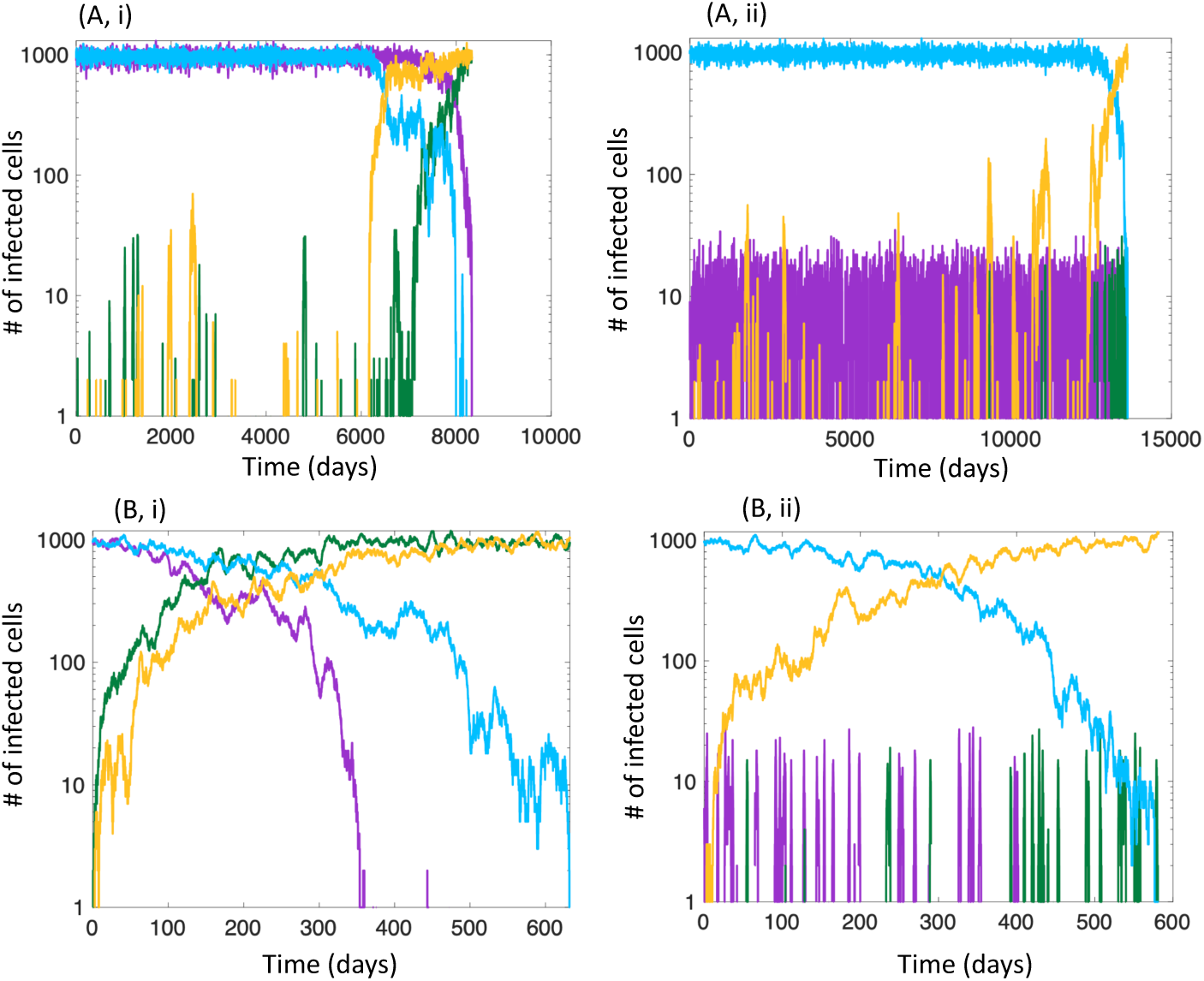
Individual realizations of evolutionary dynamics, based on Gillespie simulations of model (1). (A) Relatively low mutation rate, *μ=2×10^-5^*. (B) Higher mutation rate, *μ=2×10^-3^*. Panels (i) assume a weak CTL response, *c=0.*0001. Panels (ii) assume a strong CTL response, *c=10*. Other parameters are identical to Fig 2. Blue and purple indicate wild-type virus in the F and EF compartments, respectively. Yellow and green indicate mutant virus in the F and EF compartments, respectively.

We have so far discussed results in the context of a specific set of parameters. Variations in key parameters are explored in the Supplementary Materials. Variations in the fitness of the advantageous mutants leads to the same trends, as long as the degree of the fitness advantage is not too large (Figure S4A). The infected cell population size within the lymphoid compartments is a source of uncertainty in the model. We repeated our simulations assuming different equilibrium population sizes of infected cells (and simultaneously adjusting the mutation rate per infection to keep the total rate of mutant production constant). Again, trends remain qualitatively the same; they become more pronounced for larger populations (Figure S4B). While we have chosen parameters such that the basic reproductive ratio of the virus, R_0_≈8, variations in this measure have been observed in clinical data [43]. In Figure S4C, we show that trends remain robust for smaller and larger values of R_0_, with the compartmental structure having an increasingly large effect for higher values of *R_0_*. Another source of uncertainty concerns the rate at which populations move between compartments.

Increasing the migration rates over a range in which pronounced compartmentalization of virus load is still observed for strong CTL responses, the observations remain robust (Figure S4D).

## Discussion and Conclusion

Using mathematical models, we have shown that the compartmental structure of the secondary lymphoid tissues can influence the *in vivo* evolutionary dynamics of HIV-1 mutants that are characterized by a low to moderate fitness advantage compared to the wild-type virus. This is seen in different kinds of evolutionary measures: (i) Compartmental structure can lead to weakened selection and to a non-monotonic dependence of the mutant fixation probability on the strength of the CTL response, driven mainly by differences in infected cell population sizes between the two compartments. (ii) Compartmental structured delays the conditional fixation times of mutants. (iii) Compartmental structure slows down the rate of mutant generation, and this becomes more pronounced for stronger CTL responses. (iv) Compartmental structure delays the overall time of mutant invasion, and this again becomes more pronounced for stronger CTL responses.

The degree of virus compartmentalization has been shown to increase for stronger CTL responses in SIV-infected macaques [22, 30]. Indeed, the highest degree of virus compartmentalization has been found in so called elite controllers. In such animals, the CTL response is thought to clear productive virus from the extrafollicular compartment, and the small number of infected cells observed in the EF compartment are thought to arise due to recent immigration from the follicular compartment, where the virus can persist unopposed by CTL[22]. In such a setting of elite control, our modeling results suggest that the invasion of advantageous mutants is delayed the most compared to a hypothetical setting where the virus replicates at the same overall abundance but without compartmental structure. Because virus evolution is thought to contribute to progression of disease, this suggests that the compartmental structure in the secondary lymphoid tissue can lead to a slower rate of disease progression than expected from virus evolution models that do not take into account the compartmental architecture. In other words, while the immune-privileged nature of the follicular compartments contributes to persistent virus replication and continued virus evolution in the face of efficient antiviral CTL responses, the rate of virus evolution is predicted to proceed slower compared to a setting in which the same virus load existed in a more mixed system. The immune-privileged nature of the follicular compartment might thus be less of a driver of disease progression than previously thought, even though it contributes to persistent virus replication.

There remain uncertainties regarding several parameter values of the model, as described in the Results section. Our exploration of the parameter space with computer simulations (Supplementary Materials), however, provides confidence that our results hold broadly for mutants with a low to intermediate fitness advantage (which is biologically probably most relevant). Details of the patterns can change, e.g. if the mutation rate relative to the population size is varied, but the conclusion remains robust that virus evolution is expected to proceed more slowly in the context of the assumed compartmental structure compared to a system with equivalent virus load but lack of compartmental structure. There are also uncertainties with respect to the structure of the mathematical model, in particular the way in which CTL responses have been modeled. We assumed that virus-specific CTL expand in response to antigenic stimulation, but this can be expressed mathematically in different ways. We chose a formulation in which the dynamics are more stable, where the rate of CTL expansion is not proportional to the number of CTL. If the rate of CTL expansion was assumed to be proportional to the number of CTL, this would be more similar to a predator-prey type model with more extensive population fluctuations, which could have an impact on how the mutant fixation probability depends on the strength of the CTL response. It is not clear whether such a model would be more realistic, and this is also more difficult to study with stochastic simulations because the population oscillations can lead to extinctions events in such settings. It is important to keep in mind, however, that CTL dynamics can be formulated in a variety of different ways that all capture biologically realistic assumptions, and that this might impact results.

This work adds to the notion that the compartmental structure of the secondary lymphoid tissue can have substantial impacts on the *in vivo* evolutionary dynamics of HIV. In previous work, we have established this specifically for the case of CTL escape mutant evolution [33], and in the current study have extended it more generally to the emergence of advantageous mutants that are not related to CTL recognition. These insights have important practical implications. Mathematical models have been a very useful tool for studying the *in vivo* dynamics and evolution of HIV-1 infection. They can be informed by data, such as the overall virus load in SIV-infected macaques or in PLWH. We have shown that when it comes to studying the timing of *in vivo* evolutionary processes and hence disease progression, mathematical models that do not take into account the compartmental structure of the secondary lymphoid tissue might lead to inaccurate and misleading results. Assuming the same overall virus load, we have demonstrated that models with compartmental structure predict substantially slower rates of virus evolution compared to models that lack this compartmental tissue architecture.

## Supporting information

Supplementary details about the mathematical models

## Notes

### Competing Interest Statement

The authors have declared no competing interest.

