## Supplementary details about the mathematical models for "Compartmental structure in the secondary lymphoid tissue can slow down *in vivo* HIV-1 evolution in the presence of strong CTL responses"

### Supplementary Information

#### 1. Migration rates and compartmentalization in model (1), main text

Figure S1A and B show how the equilibrium number of infected cells and CTL in both compartments changes with an increase in the infected cell migration rate, $\eta$ (Figure S1A), and the rate of CTL movement from the EF to the F compartment, *g* (Figure S1B), assuming a

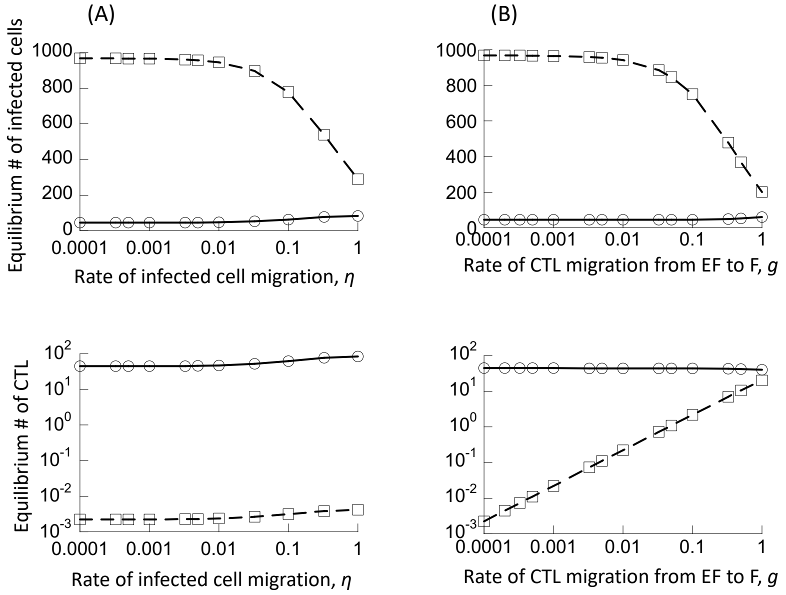


**Figure S1**. Basic properties of model (1) without mutant evolution ($\mu$=0). The equilibrium number of infected cells (top row) and the equilibrium number of CTL (bottom row) are shown as a function of select model parameters. (A) Dependence on the migration rate of infected cells between compartments, $\eta$. (B) Dependence on the migration rate of CTL from the EF to the F compartment, *g*. Dashed lines show the populations in the F compartment, solid line in the EF compartment. Base-line parameters are given as follows. $\lambda_f$=500; $\lambda_e$=500; $\delta$=0.01; *a*=0.45; $\beta_e=\beta_f=7\text{x}10^{-5}$; *p*=0.05; *b*=1; *c*=1; $\eta$=0.0001; *g*=0.0001; *h*=1.

relatively strong CTL response. As these migration rates are increased from low to high, we see that the equilibrium number of infected cells in the F compartment declines continuously, first slowly, and then faster for larger migration rates. The more pronounced decrease of infected cells at steady state in the F compartment for larger migration rates comes about because of a larger degree of mixing between the two compartments, which exposes an increasing number of infected cells from the follicular compartment to CTL-mediated activity. This is not observed in data from SIV-infected macaques, and therefore these higher migration rates are not biologically relevant. A range of lower migration rates, however, is compatible with the lack of

CTL-mediated control in the F compartment (Figure S1A and B), because in this range, increasing the migration rate results in a minimal reduction in the equilibrium number of infected cells in the F compartment. As a baseline, we will assume that the migration rates are at the lower end, i.e. $\eta = g = 10^{-4}$, and perform the initial analysis with these values. We subsequently investigate how variation in the parameters $\eta$ and $g$ (within ranges compatible with experimental data) influences the outcomes in the model.

#### 2. The one-compartment control model

The main text defines a compartmental model of virus dynamics to study the evolution of advantageous mutants by human immunodeficiency virus (HIV). In particular, we distinguish between the follicular and extrafollicular compartments in the secondary lymphoid tissues. The results obtained for these models are compared to those seen in a corresponding single-compartment control model, which is outlined as follows.

We denote the populations of uninfected cells, infected cells, and CTL by *x*, *y*, and *z*, respectively. The time evolution of these populations is given by the following set of ordinary differential equations:

$$\frac{dX}{dt} = \lambda - \delta X - \beta_1 XY,$$

$$\frac{dY_1}{dt} = \beta_1 XY_1 - aY_1 - pY_1 Z,$$

$$\frac{dZ}{dt} = cY_1 - bZ.$$

Similar models have been widely used in the literature, e.g. [1-4]. Target cells are produced with a rate $\lambda$, die with a rate $dX$, and become infected by virus with a rate $\beta_1 XY_1$ (assuming that virus is in a quasi-steady state). Infected cells are characterized by a basic death rate $aY_1$, and are killed further by CTL with a rate $pY_1Z$. The CTL population expands with a rate $cY_1$ and dies with a rate $bZ$.

In the absence of the CTL, the virus population establishes a persistent infection if its basic reproductive ratio $R_0 = \beta_1 \lambda / da > 1$. The system then converges to a stable equilibrium, given by $X^{(1)} = a/\beta_1$, $Y_1^{(1)} = \lambda/a - \delta/\beta_1$, $Z^{(1)} = 0$. If $c > 0$, the CTL response expands in the presence of the infection. In this case, the populations converge towards the following stable equilibrium.

$$X^{(2)} = \frac{a\, b\, \beta_1 - c\, \delta\, p + \sqrt{(a\, b\, \beta_1 - c\, \delta\, p\,)^2 + 4b\beta_1^2 \lambda cp}}{2b\beta^2},$$

$Y_1^{(2)} = \frac{b(\beta_1 X^{(2)} - a)}{cp}$,

$Z^{(2)} = \frac{\beta_1 X^{(2)} - a}{p}$.

Next, we incorporate an advantageous mutant virus strain into the model, described by the subscript “2”; the infected cells are hence denoted by $Y_2$. The modified ODEs are given as follows:

$\frac{dX}{dt} = \lambda - \delta X - \beta_1 X Y_1 - \beta_2 X Y_2$,

$\frac{dY_1}{dt} = \beta_1 X Y_1 (1 - \mu) - a Y_1 - p Y_1 Z$,

$\frac{dY_2}{dt} = \mu \beta X Y_2 + \beta_2 X Y_2 - a Y_2 - p Y_2 Z$,

$\frac{dZ}{dt} = c(Y_1 + Y_2) - bZ$.

The mutant virus strain is characterized by its specific rate of infection, $\beta_2$, which is assumed to be larger than the value of $\beta_1$, thus representing an advantageous mutant. It is produced during infection events with a probability $\mu$. We assume that the $R_0$ of both virus strains is greater than unity, and that the CTL population expands. Since the mutant is advantageous, it will invade and exclude the wild-type infected cell population. The system thus converges to the following equilibrium

$X^{(3)} = \frac{a\, b\, \beta_2 - c\, \delta\, p + \sqrt{(a\, b\, \beta_2 - c\, \delta\, p\,)^2 + 4b\beta_2^2 \lambda cp}}{2b\beta^2}$,

$Y_1^{(3)} = 0$,

$Y_2^{(3)} = \frac{b(\beta_2 X^{(3)} - a)}{cp}$,

$Z^{(3)} = \frac{\beta_2 X^{(3)} - a}{p}$.

#### 3. Fixation probability in a two-patch Moran process model with unequal patch sizes and migration

We implemented a constant population death-birth Moran process model with migration, considering two patches with population sizes that are identical to the equilibrium infected cell population sizes in the F and EF compartments in model (1) for the various values of *c* (immune responsiveness). At each iteration, the following events can occur in the simulation. With a probability $1-p_{mig}$, a death-birth event occurs within a patch. A cell in either patch (wt or mutant in patch 1 or 2) is selected randomly with a probability given by the fraction of that cell population among all cells in both patches. The selected cell is removed (death event). Then one of the remaining cells in the same patch is chosen and replaces the removed cell through a birth event. The cell for the birth event (either wt or mutant) is chosen with a probability proportional to the fitness of that cell. Specifically, if we denote the number of wild-type and mutant cells in a given patch by $N_1$ and $N_2$, respectively, a mutant is picked with a probability $(1+s)N_2/ [N_1+(1+s)N_2]$, where *s* denotes the fitness advantage of the mutant. With a probability $p_{mig}$, a migration event occurs. Again, a cell is chosen randomly from all cells in both patches, with a probability given by the fraction of that cell type among all cells. This cell then replaces a randomly chosen cell in the other patch, which is a migration event. Because this migration reduces the population size in the current patch by one cell, the migrating cell is replaced by a division event in the current patch, based on the fitness of the wild-type and mutant cells, as described above. Migration is thus coupled to a division event to ensure a constant population.

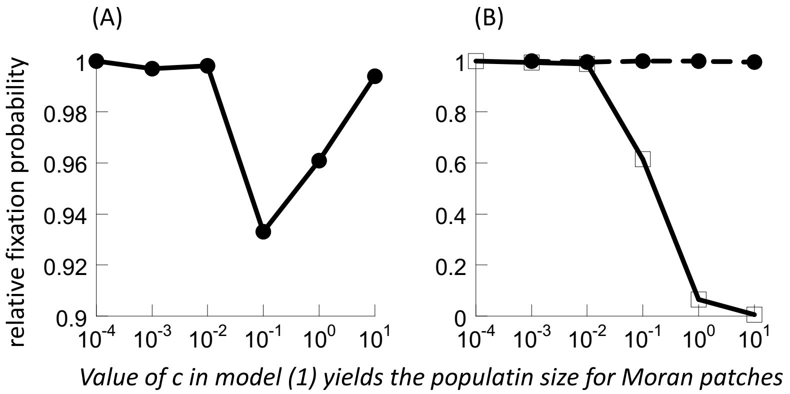


**Figure S2**. Numerically obtained relative fixation probability in the Moran process simulation with two patches and migration (normalized against the standard Moran process prediction for undivided populations). (A) Mutant is placed randomly in either patch, with a probability determined by the relative population size in each patch. (B) Mutant is always placed in the larger (dashed line) or the smaller (solid line) patch. For ease of comparison, the x-axis shows the value of the parameter c in model (1) from the main text that yields the patch population sizes assumed in the Moran process simulations. The simulations assumed a migration probability $p_{mig}$=0.0001, and a 1% mutant advantage. Fixation probabilities are based on 10,120,000 realizations of the simulation for each point.

At the beginning of the simulation, all cells are wild-type, and one cell is randomly chosen to be replaced by a mutant. This can occur in either patch, with a probability given by the fraction of the cells in a given patch. The simulation is then run until the mutant either goes extinct or reaches fixation in both patches. The fixation probability is determined numerically based on repeated realizations of the simulation, as described in the main text. Results are shown in Figure S2A. Different populations sizes in the patches are considered, corresponding to those in model (1) in the main text for specific immune response strengths (parameter *c*). The same non-monotonic dependence of the fixation probability on "c" is observed as seen in Figure 2A in the main text. The weakened selection is the result of those realizations, where the initial mutant is placed into the smaller patch. In Figure S2B, simulation results are presented when the mutant is always placed either into the smaller or into the larger patch. When the mutant is always placed into the smaller patch, we see monotonically reduced selection for more unequal population sizes among the patches (higher value of c in model (1), main text). No weakened selection is observed when the mutant is always placed into the larger patch.

#### 4. Effect of population size and fitness on mutant fixation probabilities

Here, we show how the mutant fixation probability in the two-compartment model (shown in Figure 2A in the main text) depends on the population size and mutant fitness. This is shown in Figure S3. The weakened selection for intermediate values of *c* becomes less pronounced for

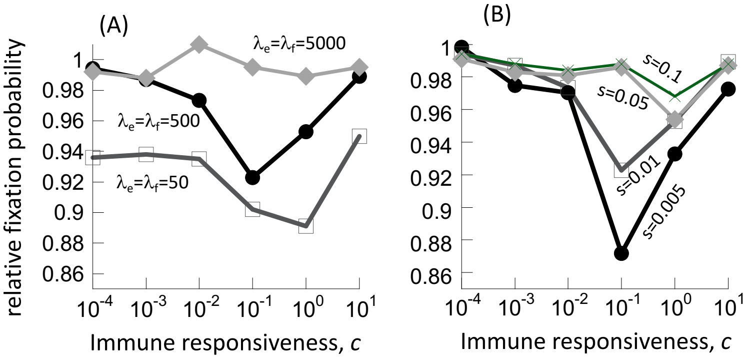


**Figure S3**. Relative mutant fixation probabilities in the two-compartment model as a function of the immune responsiveness *c* for (A) different population sizes and (B) degrees of mutant fitness (compare to Figure 2A in the main text, same parameters unless indicated otherwise). In the model, the infected cell population size is directly proportional to the parameters $\lambda_e$ and $\lambda_f$, and the values are indicated in the graphs. For $\lambda_e=\lambda_f=500$, $\beta_1=0.00007$. We assumed that an increase/decrease in $\lambda$ values was associated with a proportional decrease/increase in the value of $\beta_1$, such that the basic reproductive ratio of the virus was constant. For B, we assumed $\lambda_e=\lambda_f=500$, $\beta_1=0.00007$.

larger infected cell population sizes and eventually vanishes. The same applies to larger mutant fitness advantages.

#### 5. Dependence of the average mutant fixation time on parameters

In the main text, simulations were explored where the wild-type virus population at equilibrium gives rise to advantageous mutant strains, and the average time to mutant fixation (in both the F and EF compartments) was determined numerically as a function of CTL response strength. This was done for a particular parameter set. Here, we explore how the results depend on changes to parameters. Figure S4 plots the fold increase in the average mutant fixation time in the two-compartment compared to the corresponding one-compartment control model, assuming variation in parameters. The simulations in the main text assumed a 1% fitness advantage, and this is re-plotted in the black line (with circles) in Figure S4A. In addition, simulation results are plotted that assume a 0.5% (light grey line) and 10% (dark grey line) advantage. The fold increase in mutant fixation time brought about by compartmentalization remains robust across a range of fitness advantages (Figure S4A).

The effect of total equilibrium population size is explored in Figure S4B. The population size assumed in the main text is shown in black, and 10-fold smaller and larger population sizes are shown in light and dark grey, respectively. A difference is seen for weak CTL responses, when the equilibrium population sizes of infected cells in the two compartments is similar. For smaller populations, the average fixation time is substantially longer for the subdivided population compared to the one-compartment control model. For larger equilibrium population sizes, this difference vanishes, and population subdivision into two approximately equal compartment sizes does not substantially delay mutant fixation. For strong CTL responses, however, mutant invasion is delayed to a similar and substantial extent regardless of the population sizes.

Simulations with different values of the basic reproductive ratio of the virus, $R_0$, are shown in Figure S4C. The black line shows the simulation results from the main text, where we assumed $R_0 \approx 8$, which corresponds to the average estimate in PLWH [5]. Since there is a spread of estimated $R_0$ values among different individuals, we also ran simulations with a lower (light grey) and higher (dark grey) value of $R_0$. Results remain generally robust but become less pronounced at the lower end of estimated $R_0$ values.

Finally, we varied the migration rates of infected cells between compartments, and the movement rate of CTL from the EF to the F compartment (Figure S4D). In the main text, we assumed relatively low values (black line in Figure S4D). Here, we increased the migration rate within a range in which substantial differences in infected cell population sizes between the EF and F compartment still exist for efficient CTL responses (see Figure S1 for this range). Simulation results were found to remain robust within this range of migration rates (Figure S4D).

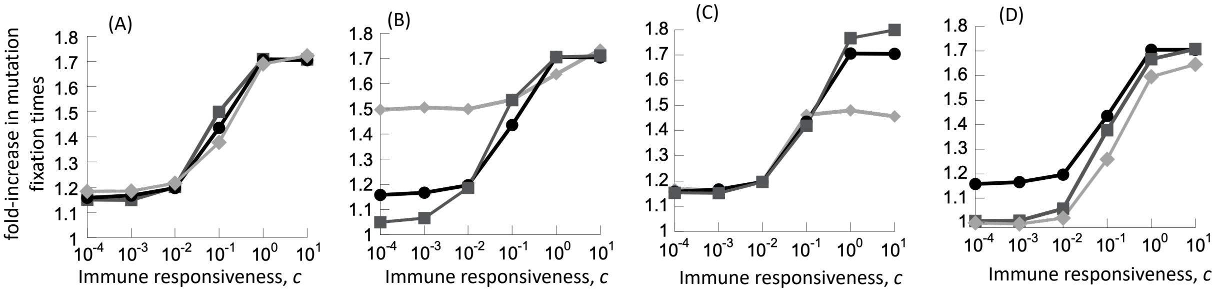


**Figure S4**. Fold increase of mutant fixation time in the two-compartment model compared to the one-compartment control model. This is based on stochastic simulations of model (1) and the corresponding one-compartment control model. Details are given in Figure 4 of the main text. The black line is always the same as presented in the main text for a particular parameter set specified in Fig 2 of the main text. We assumed a mutation rate of $\mu=2x10^{-5}$. (A) Variation in mutant fitness. s=0.005 for the light grey line, and s=0.1 for the dark grey line. (B) Variation in the population size in the two compartments. For lower population size (light grey line), we assumed For $\lambda_e=\lambda_f=50$, $\beta_1=0.007$ to keep $R_0$ constant, and $\mu=2x10^{-4}$ to keep the rate of mutant generation constant. For higher population size (dark grey), we assumed $\lambda_e=\lambda_f=500$, $\beta_1=0.00007$, and $\mu=2x10^{-6}$. (C) Variation in the basic reproductive ratio of the virus. $R_0=3$ for the light grey line, and $R_0=15$ for the dark grey line. (D) Variation in the migration rates $\eta$ and $g$. For the dark grey line, $\eta=0.01$; g=0.01. For the light grey line, $\eta=0.1$; g=0.1.
